# Information-dependent eye-hand coordination emerges from active vision

**DOI:** 10.64898/2026.05.29.726887

**Authors:** Jingwen Zhao, Dorian Verdel, Ying Tan, Etienne Burdet

## Abstract

In daily activities, humans rely on visual information to plan hand movements, making the extraction of task-relevant information through eye gaze a key aspect of motor control. Behavioral studies have revealed characteristic saccade-pursuit patterns, likely governed by shared neural circuits, which enable an efficient reduction of task-related uncertainty.

However, a unifying computational principle explaining the emergence of these patterns in continuous tasks such as reading or driving is still lacking. Here we propose a dual stochastic model predictive control formulation of active vision, in which eye movements are continuously controlled to minimize task-relevant uncertainty and build an internal model used for hand movement planning. Through experiments manipulating the amount, density, and difficulty of future visual information, we show how eye movement patterns adapt to the information context while maintaining an invariant extraction horizon. A saccade-pursuit pattern naturally emerges from the model, which accurately predicts both eye and hand movement features observed in experiments. These results provide a principled framework for understanding the continuous regulation of human eye movements and open new perspectives for applications in robotic assistance and active perception.

## Introduction

Daily activities such as assembling a device, catching a fly, or driving in traffic rely on seamless coordination between our eyes, which gather information, and our hands, which execute the task. The principles underlying hand movements control have been extensively studied ***Milner (1992***); ***Uno et al. (1989***); ***Harris and Wolpert (1998***); ***Franklin et al. (2007***); ***Wang et al. (2016***); ***Berret et al. (2024***), providing strong evidence that motion is planned over a future horizon to counteract disturbances and uncertainty. In contrast, our understanding of the principles governing eye movement control, and its coordination with hand movements, remains limited. Behavioral studies suggest that gaze is directed to support decision-making and motor planning. For instance, during interception, gaze anticipates the predicted bounce point ***Land and McLeod (2000***); ***Arthur et al. (2024***); ***Trevi and Márquez (2025***), and when viewing tools, it is preferably drawn to its functional regions (e.g., the hammer head) rather than to manipulative parts (e.g., the handle) ***Belardinelli et al. (2016***); ***Pilacinski et al. (2021***). In continuous tracking tasks, efficient information gathering is essential for planning over a future horizon ***Bashford et al. (2022***), as illustrated by experienced drivers who consistently look farther ahead of their vehicle than novices (***Figure 1A) Xu et al. (2022***). In the present paper, we propose that such behaviors emerge from an *active vision* principle, whereby the central nervous system (CNS) controls gaze to acquire task-relevant information and enable efficient movement with minimal effort.

**Figure 1.**
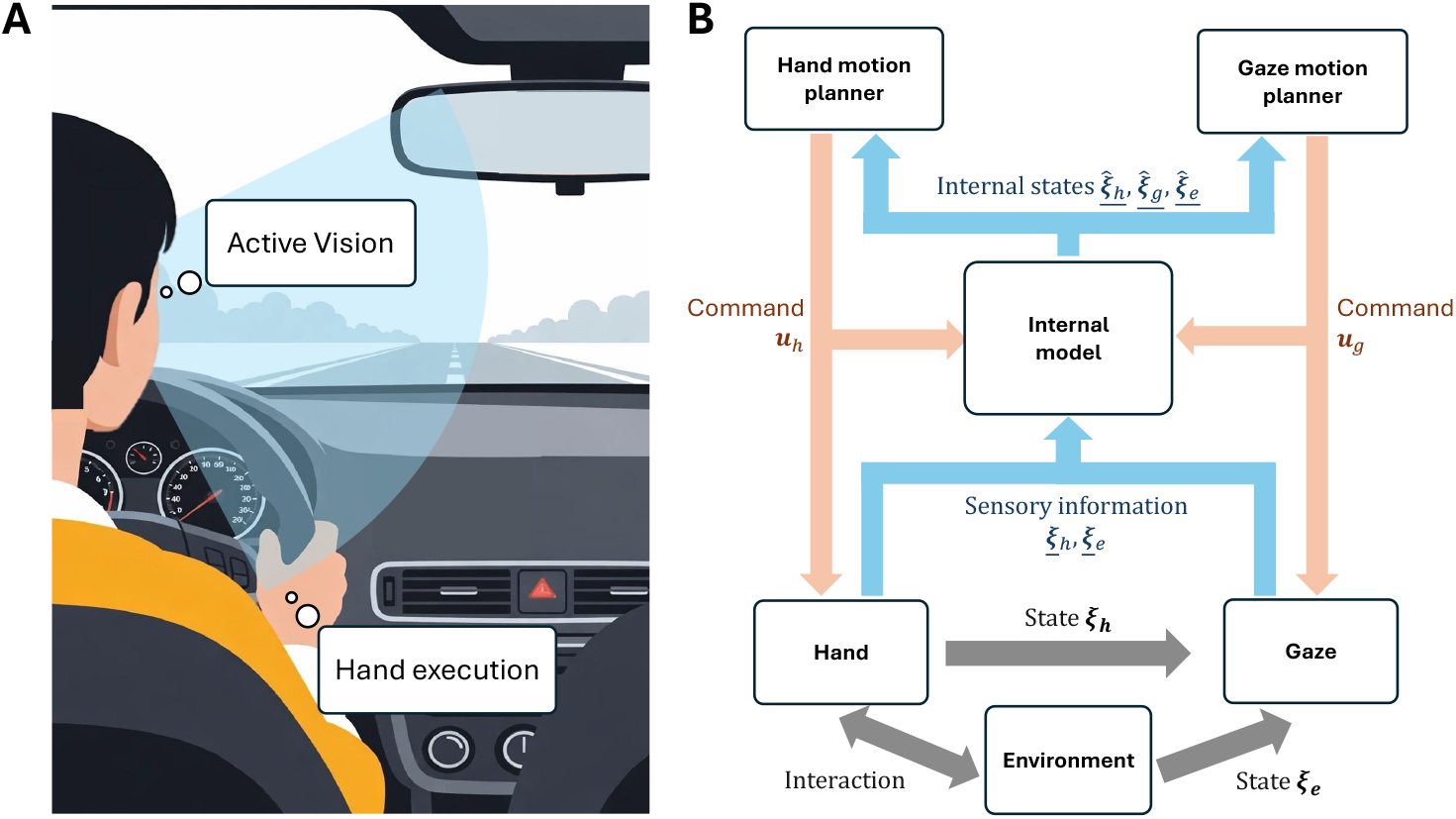
Active vision for eye-hand coordination. **A**: Driving exemplifies a continuous eye-hand coordination task requiring active vision with coordinated hand movements. Eye movements are planned to continuously acquire information (i.e., reduce uncertainty) about the road, the vehicle, and the hands, while the hands act on the steering wheel to control the vehicle based on visual input. **B**: Implementation of active vision using a dual model predictive control (MPC) framework. Noisy observations of the environment ξ_*e*_ and hand ξ_*h*_ are acquired through gaze. These observations, together with efferent copies of eye and hand motor commands and past observations, are integrated via a state estimator to update the internal model of the task. This yields (i) estimates of the current states of the environment 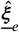, eye gaze 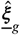, and joint state of the hand 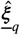, and (ii) predictions of how gaze movements affect task and hand uncertainty. These estimates are then used to compute optimal eye and hand control inputs **u**_*g*_ and **u**_*h*_ over a finite time horizon to interact with the environment.

This principle must consider the characteristics of the eyes as sensors. In humans (as well as haplorhine primates, birds of prey, and some reptiles and fish), visual acuity peaks at the fovea and declines towards the periphery ***Curcio et al. (1987***); ***Stewart et al. (2020***). Although the eyes have low inertia ***Robinson (1964***), their precise control by three antagonist muscle pairs entails non-negligible effort and is often coordinated with head and body movements ***Guitton and Volle (1987***). Together with neurophysiological constraints ***Schütz and Stewart (2025***), this leads to characteristic patterns of saccades and pursuit (or fixation) ***Robinson (1964***); ***Ko et al. (2010***); ***Danion and Flana- gan (2018***); ***Mathew et al. (2019***). A computational account of active vision should naturally give rise to these patterns.

Existing models typically treat saccadic and pursuit movements separately, often relying on predefined targets or velocity profiles. Approaches include linear feedback control ***Robinson (1973***); ***Krauzlis and Lisberger (1989***), minimum variance ***Harris and Wolpert (1998***), optimal control ***Zhou et al. (2018***), and stochastic optimal control ***Varsha et al. (2021***); ***Vasudevan et al. (2023***). In parallel, probabilistic models of visual attention emphasize goal dependence ***Ballard and Hayhoe (2009***), prior knowledge ***Henderson (2017***), context ***Eckstein (2017***), and visual saliency ***Walter and Bex (2022***), predicting sequences of fixation targets rather than modeling eye movements. These approaches include dynamics Bayesian networks ***Ballard and Hayhoe (2009***); ***Hayhoe and Ballard (2014***), Markov decision processes ***Ma et al. (2022***); ***Chen et al. (2021***), stochastic optimal control ***Vasilyev (2019***),information gain maximization ***Renninger et al. (2005***); ***Najemnik and Geisler (2005***); ***Paeye et al. (2016***); ***Hoppe and Rothkopf (2019***), and intrinsic reward formulations ***Xu-Wilson et al. (2009***); ***Rothkirch et al. (2013***). As a result, they typically capture either saccadic behavior or discrete fixation sequences.

Attempts to jointly model saccades and pursuit often rely on switching mechanisms based on eye-target error ***De Brouwer et al. (2002***) or its variability ***Coutinho et al. (2021***), effectively treating them as separate processes. However, neuroscience evidence indicates that the underlying premotor pathways and oculomotor subsystems are largely shared ***Liston and Krauzlis (2003***); ***de Xivry and Lefèvre (2007***); ***Goettker and Gegenfurtner (2021***). A unified stochastic optimal feedback controller has been proposed to account for saccade–pursuit coordination ***Crevecoeur and Kording (2017***), but it primarily captures motor aspects and does not address the key role of the eyes as information-gathering sensors. Moreover, humans can adust motion plan online in response to changing visual input ***Van Gisbergen et al. (1987***); ***Doré et al. (2025***), suggesting prospective, horizon-based control rather than purely reactive strategies.

Efficient eye–hand coordination is not innate but develops through sensorimotor experience, as the relationships between visual input, body state, and motor output are learned during infancy and refined through childhood ***von Hofsten (1989***); ***Corbetta and Snapp-Childs (2009***). Here we propose a model of coordinated eye and hand movement in adults based on an active vision principle. The model generates saccade-pursuit patterns from a single continuous optimization process, incorporating key neurophysiological constraints and an internal task representation to enable efficient information acquisition and action.

To evaluate the model, we recorded eye and hand movements in 17 participants performing a trajectory-tracking task in which future target motion was visible over a finite horizon, allowing participants to extract visual information about future desired hand positions. Previous work with a similar task has shown that people plan in a MPC-like fashion ***Hafs et al. (2026***), and that their movement variability decreases and planning horizon increases over practice ***Bashford et al. (2018***). We systematically manipulated three properties of available information: (i) temporal horizon (how far into the future information is available for motion planning), (ii) spatial density (how much information is contained per unit space), and (iii) the effort required for accurate tracking, to assess their influence on gaze strategies and performance.

## Results

### Active vision for eye-hand coordination

We developed an active eye-hand control strategy according to which eye movements are viewed as selecting gaze positions to acquire task-relevant information through minimizing task-relevant uncertainty. We modeled the continuous coordination of eye and hand movements as a dual model predictive control (MPC) system, as schematized in ***Figure 1***B. This system comprises three key elements: (i) a gaze motion planning control strategy, (ii) an internal model that estimates the current state of the environment and hand, and (iii) a hand motion planner that uses this knowledge to generate motor commands. Our model describes how the gaze ξ_*g*_ and hand ξ_*h*_ movements resulting from the motor commands ***u***_*g*_ and ***u***_*h*_ depend on the environment state ξ_*e*_ in task space, e.g. to track a target trajectory. Mathematical notation follows standard conventions: bold symbols denote vectors (e.g. **v**), bold capital letters denote matrices (e.g. **M**) and underlined symbols denote stochastic variables (e.g. *v*). Details on the implementation are provided in the Methods.

To simulate eye gaze behavior, we first assume that the eye dynamics

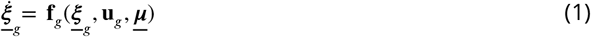

where the nonlinear mapping **f**_*g*_ is assumed to be twice differentiable an depends on the eyes gaze state ξ (including position and velocity), the time-varying gaze motor command **u**_*g*_ and motor noise **μ**. The active control of these eye dynamics must account for both environmental and internal sources of uncertainty, which can stem from multiple sources. Environmental uncertainty *U*_*e*_ relates to the target uncertainty in task space and its physical properties, while internal uncertainty *U* _*h*_ arises from sensorimotor noise affecting the estimation of the hand state.

The objective of gaze control at a given time *t*_*c*_ is expressed through the cost

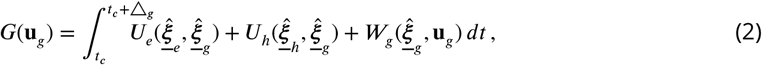

over an information gathering horizon △_*g*_, where 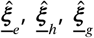 denote the time-varying estimated states of the environment, hand, and eye gaze, respectively. Importantly, uncertainties *U*_*e*_, *U*_*h*_ depend on the visual information gained with current gaze state ξ_*g*_. Minimizing this cost trades off uncertainty reduction in the environment and hand movement against the energetic cost of eye movements 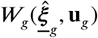.

With the high density of cone cells in the human fovea, high-acuity vision is restricted to a small region around the optical center, with acuity decreasing toward the periphery. We model this by assuming that visual noise increases quadratically with retinal eccentricity, consistent with cone cells density distribution ***Curcio et al. (1987***). For a point [*x, y*]^′^ in the visual field relative to the gaze [*x*_*g*_, *y*_*g*_]^′^, the noisy observation is

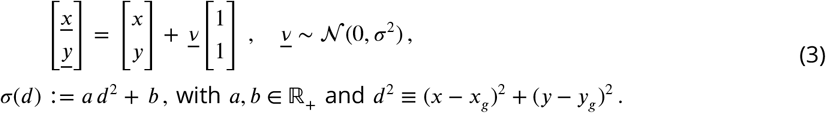

Consistent with reported hand reaction times ***Bekkering et al. (1994***); ***Dean et al. (2011***); ***Verdel et al. (2026***), each observation becomes available for hand movement planning after a delay of 200 ms. The CNS is assumed to integrate current and past observations optimally using a state estimator. The uncertainties *U*_*e*_ and *U*_*h*_ are computed from the covariance matrices of the estimated environment and hand states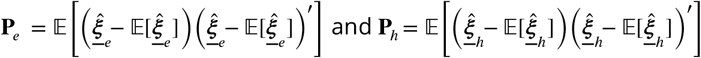,respectively.

Hand dynamics are modeled as:

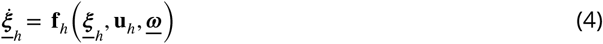

where the nonlinear mapping **f**_*h*_ is assumed to be twice differentiable, and ξ_*h*_ and **u**_*h*_ denote the hand’s state and control inputs (represented in task or joint space), and ω_*h*_represents motor noise. Hand movements are planned over a receding horizon △_*h*_ by minimizing

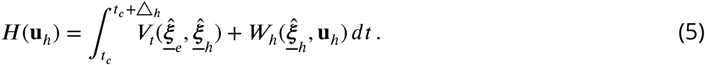

where *V*_*t*_ captures task performance and *W*_*h*_ reflects energetic cost, both function of time-varying state and control variables.

Together, ***Equations 1–5*** define a unified framework for eye-hand coordination that can accommodate a broad range range of tasks, state representations, and cost structures. The cost terms must satisfy *V*_*t*_ ≥ 0 and *W*_*h*_ > 0, allowing for a broad range of formulations, including standard quadratic costs ***Todorov (2005***), mechanical work ***Berret et al. (2008***); ***Gaveau et al. (2021***); ***Verdel et al. (2023a***), and the incorporation of additional factors such as reward ***Shadmehr et al. (2010***); ***Rigoux and Guigon (2012***) or time ***Berret and Jean (2016***); ***Verdel et al. (2023b***).

### Model validation

To validate the model, we simulated a trajectory tracking task requiring continuous gaze movements to sample task-relevant visual information, and continuous hand movements to execute motor responses (***Figure 2***A). A target trajectory was presented within a sliding temporal window and had to be tracked using horizontal wrist flexion-extension movements. Simulations used physiological parameters representative of an average human from the literature, while systematically varying viewing horizon, motion frequency, and scrolling velocity to modulate the amount, density, and quality of visual information. Overall, both the model prediction and the experimental result shows that the gaze behavior adapts to different information conditions while maintaining a relatively consistent information-gathering horizon. We proceed to exam the results by examining the local motion pattern within a trial and by comparing the average behavior across different information conditions.

**Figure 2.**
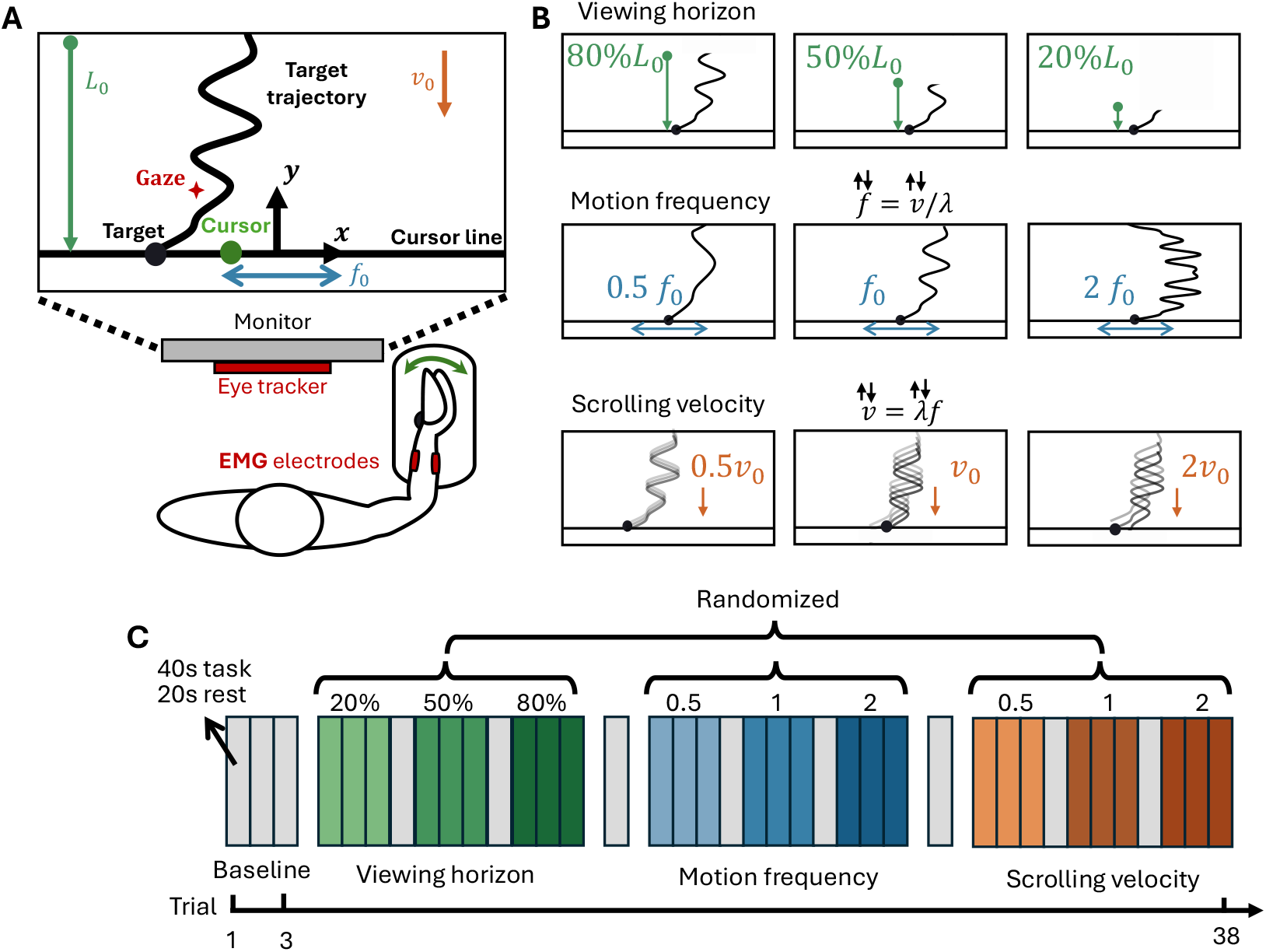
Setup and experiment to investigate active vision. **A**: Illustration of the trajectory tracking task. A future desired trajectory is displayed over a viewing horizon *L*_0_ = 26.8 cm and scrolls down at velocity *v*_0_ = 13.4 cm/s, corresponding to a wrist movement frequency *f*_0_ = 0.1 Hz. In each trial, the participant tracked the trajectory for 40 s. **B**: Task parameters were varied to probe different information and task difficulty conditions: (i) viewing horizon *L* ∈ {80%*L*_0_, 50%*L*_0_, 20%*L*_0_}, (ii) wrist movement frequency *f* ∈ {0.5*f*_0_, *f*_0_, 2*f*_0_}, and (iii) scrolling velocity *v* ∈ {0.5*v*_0_, *v*_0_, 2*v*_0_}. **C**: Experimental protocol. The session began with three baseline trials, followed by three blocks presented in a random order (one per task variation), each containing three consecutive trials per variation followed by a washout, resulting in eleven trials, presented in random order. Blocks were performed in random order and separated by washout trials, which were identical to baseline trials.

The model naturally produced characteristic saccade–pursuit patterns observed in human behavior ***Danion and Flanagan (2018***); ***Mathew et al. (2019***). As shown in ***Figure 3***A–D, gaze exhibited alternating saccade and pursuit phases. Simulated saccades displayed slightly skewed bell-shape velocity profiles, with peak velocities ranging from 65 ^°^/s to 170 ^°^/s and durations between 80 ms and 100 ms, consistent with reported values for movements of similar amplitude ***Van Opstal and Van Gisbergen (1987***). Specific for this task, saccades occurred primarily in the vertical direction, followed by downward pursuit along the trajectory, with some drift in the horizontal direction. The wrist motion controller successfully used the acquired visual information to achieve accurate tracking with low effort (***Figure 3***E,F).

**Figure 3.**
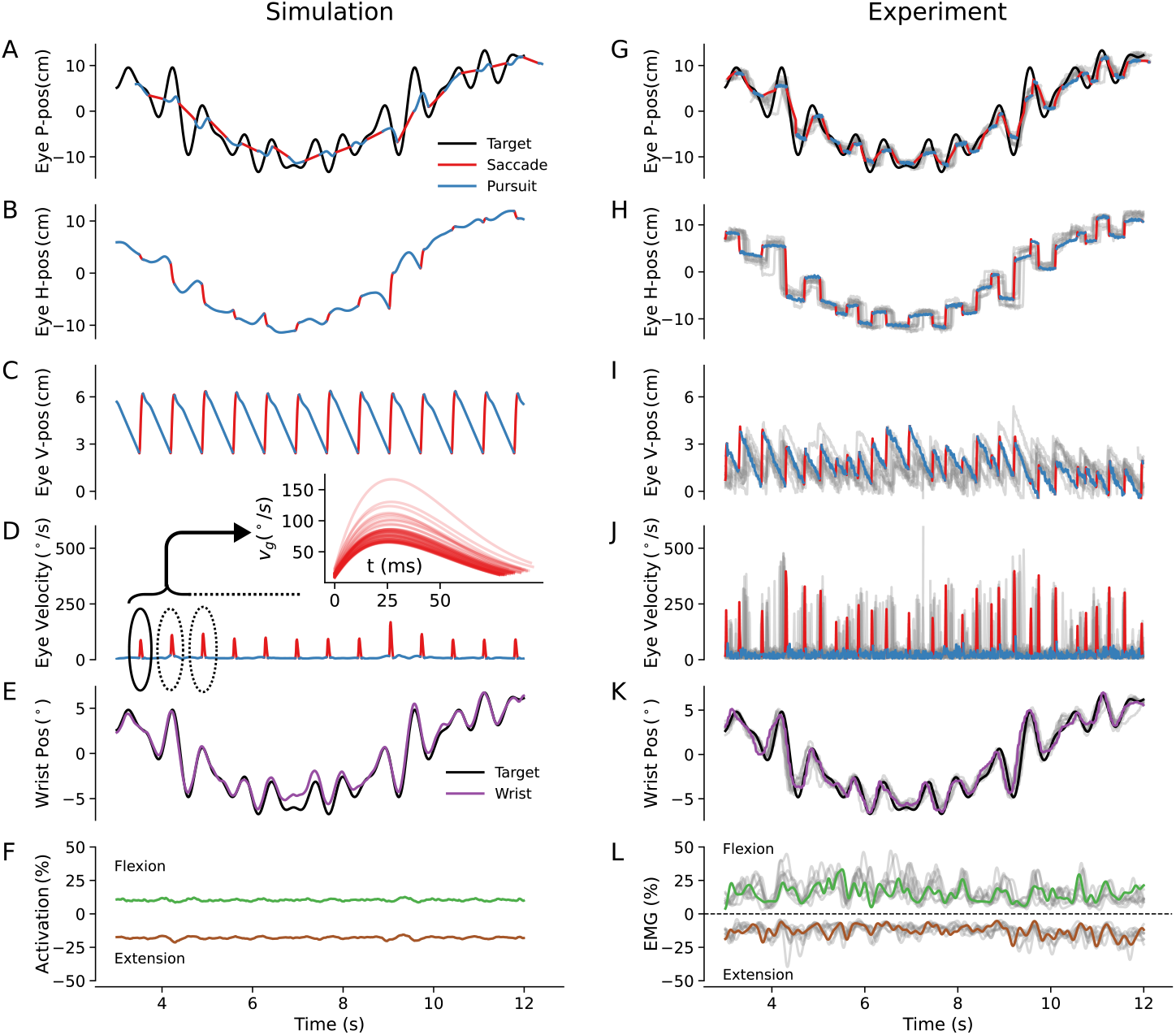
Experimental and simulated eye and wrist movement patterns. The simulation column (left) shows a segment of model output for a selected parameter set, sampled at 100 Hz. The experiment column (right) shows an example data segment from a human participant for the same target trajectory (colored), with additional repetitions of the same condition from the same participant overlaid in gray. **A,G:** Eye gaze projected onto the target trajectory in the horizontal position-time plane. **B,H:** Horizontal component of eye gaze. **C,I:** Vertical component of eye gaze. **D,J:** Eye gaze velocity. The insert in **D** shows bell-shaped saccades generated by the model at1 kHz . **E,K:** Wrist position and target trajectory. **F,L:** Flexor (positive) and extensor (negative) muscle activity; in **F**, model-predicted activations are scaled to match the participant’s average EMG level.

We further evaluated the model by varying task conditions (as described in ***Figure 2***B) and comparing simulated behavior with experimental data from 17 participants performing the same task (***Figure 2***C). Eye movements, wrist kinematics, and muscle activity were recorded.

Consistent with model predictions, participants exhibited saccade–pursuit patterns (***Figure 3***G– J). Gaze was primarily directed toward the incoming trajectory rather than the cursor, as seen in the vertical gaze position (***Figure 3***I). Tracking performance (***Figure 3***G,H) and muscle activity (***Figure 3***K,L) were comparable to model predictions, although experimental data showed greater variability.

Quantitative analyses of gaze behavior (***Figure 4***) revealed that reducing the viewing horizon decreased both gaze elevation (*V-position*) and corresponding *lead time* in the target trajectory, particularly in the 20% condition. Experimental results confirmed these trends, with significant effects for both metrics (*F*_2,28_ = 16, *p* < 0.001, η^2^ = 0.12). In detail, the 20% visible condition led to a significantly smaller V-position and lead time than the two other viewing horizons (V-position: *p* < 0.002, Cohen’s *D* > 0.67; lead time: *p* < 0.002, Cohen’s *D* > 0.67). In contrast, gaze–target distance remained largely unchanged across viewing horizon conditions, consistent with model predictions. The model predicted that increasing motion frequency would not affect V-position or lead time, but would increase horizontal gaze–target distance (***Figure 4***A–D middle panels). The experimental data confirmed the stability of V-position and lead time, despite a slight overestimation of their mean values by the model. The predicted increase in horizontal distance between gaze and the target trajectory was also supported by the data. Specifically, there was a main effect of motion frequency on both the horizontal distance (*F*_2,28_ = 320, *p* < 0.001, η^2^ = 0.74) and the overall distance (*F*_2,28_ = 11.3, *p* < 0.001, η^2^ = 0.1) between gaze and the target trajectory. In detail, the horizontal distance increased with motion frequency (*p* < 0.001, *D* > 1.61 between all pairs), while the overall distance was smaller for the highest frequency than the other frequencies (*p* < 0.005, *D* > 0.65 in both cases). Finally, the predicted slight decrease in eye-to-target distance at 2*f*_0_ was verified in the data, but no significant change was observed for the 0.5*f*_0_ condition. Overall, modulating the density of available information did not affect the average time between information gathering and movement execution, but induced a “filtering” effect, whereby gaze was directed toward intermediate regions of the trajectory rather than its peaks (see also Supplementary ***Appendix 1—figure 1***).

**Figure 4.**
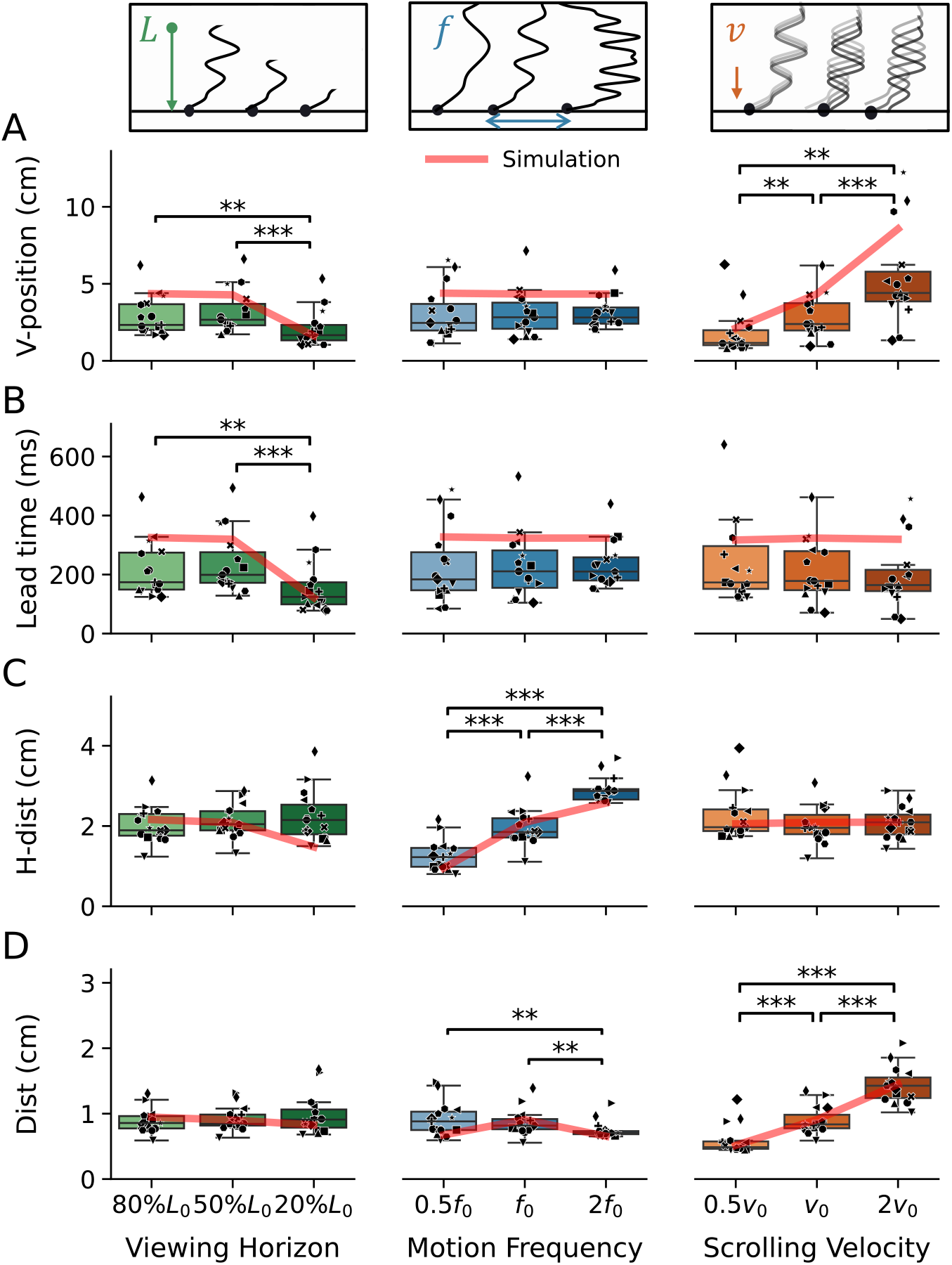
Modulation of eye gaze information-gathering patterns across conditions. In all panels, the red line indicates model predictions, and each symbol corresponds to one participant. **A:** Mean gaze elevation from the cursor line. **B:** Mean temporal lead of gaze relative to the hand cursor, computed as vertical position divided by the condition-specific scrolling velocity. **C:** Mean absolute horizontal distance between gaze and the target trajectory. **D:** Mean distance between gaze and the target trajectory.

Increasing the scrolling velocity of the target trajectory led the model to predict higher gaze positions while maintaining a constant lead time. For the distance between gaze and the target trajectory, the model predicted no change in horizontal distance but an increase in overall distance (see ***Figure 4***A–D right panels). Experimental data confirmed these predictions, revealing a main effect of scrolling velocity on both V-position and Distance (V-position: *F*_2,28_ = 15.5, *p* < 0.001, η^2^ = 0.34; Distance: *F*_2,28_ = 146, *p* < 0.001, η^2^ = 0.69). Specifically, gaze elevation increased with each increase in scrolling velocity (all *p* < 0.009, *D* > 0.66), while lead time remained unchanged. Similarly, the distance between gaze and the target trajectory increased with scrolling velocity (all *p* < 0.001, *D* > 1.37), whereas the horizontal distance did not vary significantly. Overall, higher scrolling velocities led to elevated gaze positions that preserved a nearly constant lead time of about 200 ms between visual sampling and the corresponding hand movement. The increased distance from the trajectory suggests that precise fixation becomes more costly at higher speeds. These effects were accurately captured by the model, both in terms of trends and average values.

For hand movements (***Figure 5***), the model predicted increased tracking error with a reduced viewing horizon and higher motion frequency, which was confirmed experimentally. Regarding scrolling velocity, the model predicted slightly better performance (lower RMSE) at *v*_0_ compared to other velocities. Overall, tracking performance was modulated by all conditions (viewing horizon: *F*_2,28_ = 39, *p* < 0.001, η^2^ = 0.68; motion frequency: *F*_2,28_ = 110, *p* < 0.001, η^2^ = 0.52; scrolling velocity: *F*_2,28_ = 4.0, *p* < 0.05, η^2^ = 0.78). In detail, the 20% viewing horizon condition led to an increase in RMSE (both *p* < 0.001, Cohen’s *D* > 1.49). Increasing motion frequency also resulted in larger RMSE (*p* < 0.001, *D* > 1.81 across all condition pairs), although the experiment data showed larger error at 2*f*_0_ than predicted by the model. Changes in scrolling velocity (0.5 *v*_0_ and 2 *v*_0_) led to a slight increase in RMSE compared to the *v*_0_ condition (both: *p* < 0.04, Cohen’s *D* > 0.60). In summary, reducing the viewing horizon does not impair tracking performance until the available information becomes very limited. In contrast, increasing motion frequency consistently degrades tracking performance, as it alters the desired wrist movement trajectory.

**Figure 5.**
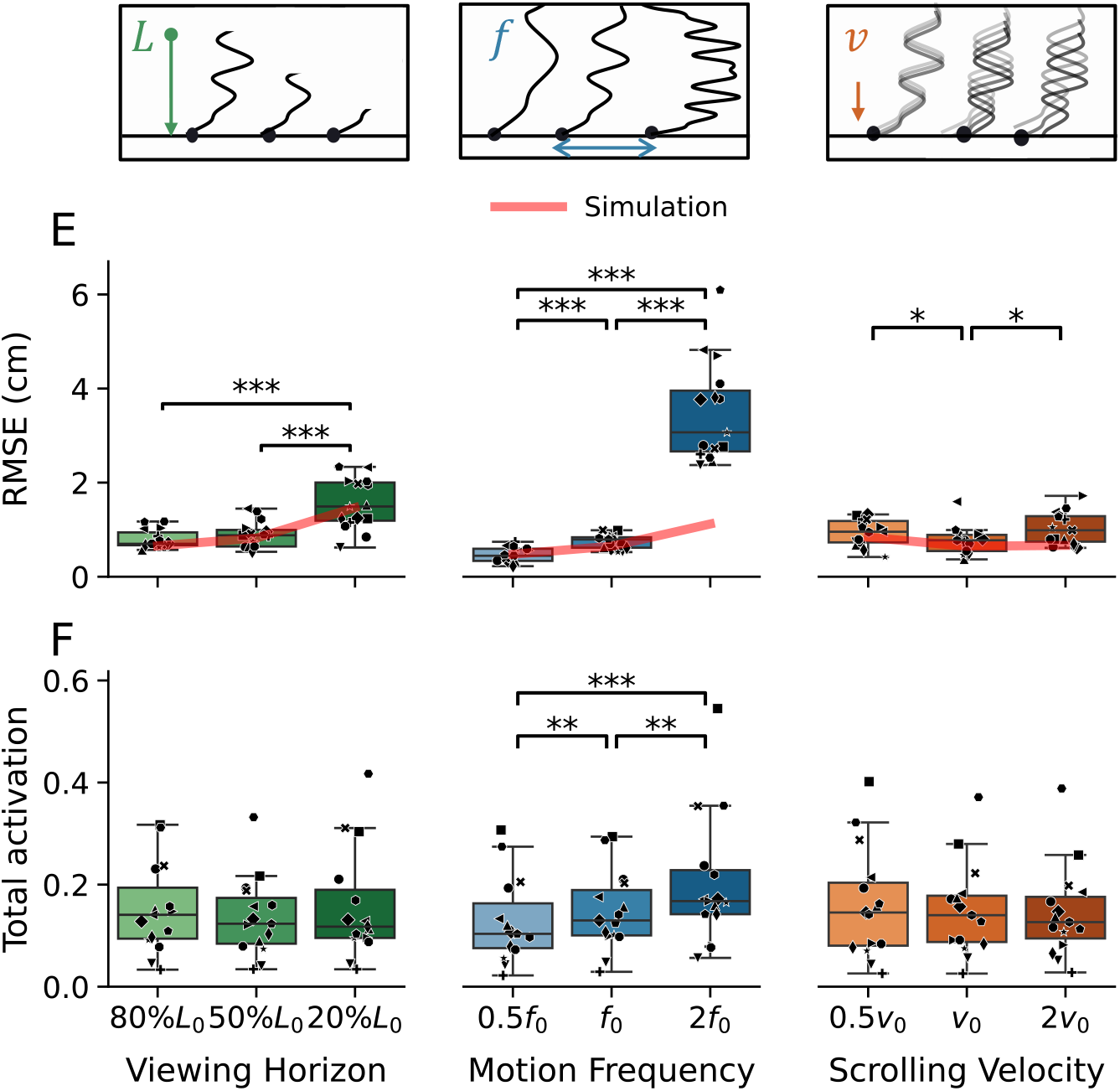
Modulation of hand movement patterns across conditions. In all panels, the red line indicates model predictions, and each symbol corresponds to one participant. **A:** Average root-mean-square horizontal distance between the hand cursor and the target trajectory. **B:** Mean total muscle activation, the flexor and extensor activity are normalized into range [0, 1] and summed.

In addition, we analyze the sum of normalized flexor and extensor activity. This *total activation* is influenced only by the motion frequency changes, and not significantly by other conditions: *F*_2,28_ = 16, *p* < 0.001, η^2^ = 0.10. Pairwise comparison shows that any increase of motion frequency induced an increase of total activation (in all cases: *p* < 0.001, *D* > 0.24). Motion frequency is also the only condition that modulated the net activation and co-contraction level as it directly changed the difficulty of the motor task.

## Discussion

In many hand–eye coordination tasks—such as hitting a ball ***Land and McLeod (2000***) or assembling a device ***Capponi et al. (2024***)—humans actively direct gaze toward task-relevant features to acquire information necessary for accurate control. Through a computational implementation of this active vision principle, our model explains how humans control eye movements to reduce task-relevant uncertainty, and use the resulting memorized information to guide hand movements. We assume that the CNS coordinates gaze to acquire informative observations while controlling the body to accomplish the task, balancing performance with the oculomotor effort for information gathering. Visual sensitivity is modeled through eccentricity-dependent noise, reflecting reduced acuity away from the fovea. Combined with internal models of the motor plant and environment, this allows prediction of future states and sensory inputs, enabling the hand to act on predicted targets despite sensory delays.

A key result of our simulations is that the active vision principle naturally generates saccade – pursuit behavior. While such patterns are well documented in tracking tasks ***Danion and Flanagan (2018***); ***Coudiere and Danion (2024***), previous models typically relied on separate controllers for saccades and pursuit ***Schütz and Stewart (2025***); ***Souto and Kerzel (2021***). Here, no explicit switching mechanism was imposed; instead, both behaviors emerge from a single objective. Intuitively, this arises from a trade-off between integrating visual information over time and maintaining the target within a high-acuity region. Smooth pursuit allows temporal integration of sensory information when retinal motion is small, improving state estimation. In general, as a target drifts away from the fovea due to motion and delays, uncertainty increases because of eccentricity-dependent noise, making corrective saccades necessary to re-center the gaze. This interplay between information accumulation and degradation provides a mechanistic explanation for the emergence of saccade–pursuit alternation.

At the sensory and motor levels, we deliberately used simplified models to limit free parameters.

For the eye, we assumed a spherical oculomotor plant with noiseless motor commands and a foveated noise profile, neglecting known asymmetries between vertical and horizontal movements ***Rottach et al. (1996***). For the hand, wrist dynamics were approximated by a third-order linear model ***Gaveau et al. (2021***), rather than a full neuromusculoskeletal representation ***Quesada et al. (2026***). Consequently, the model does not capture impedance modulation mechanisms ***Burdet et al. (2013***), which are known to respond to uncertainty and disturbances ***Franklin et al. (2008***); ***Berret et al. (2024***). However, given the absence of external perturbations and the use of constant additive motor noise, these simplifications were sufficient for capturing the dominant behaviors of interest—namely, eye movement strategies and their interaction with hand control across a broad range of information conditions.

Our model accurately predicted eye and hand movement patterns under systematic variations of the time horizon of available information, its spatial density, and the difficulty of information acquisition. Participants consistently maintained gaze approximately 200 ms ahead of the hand, consistent with compensation for sensory delays. This temporal lead was robust, producing predictable shifts in gaze position with scrolling velocity, without degrading motor performance. When less than 400 ms of future information was available, gaze patterns became constrained and shifted towards the hand, leading to a marked deterioration in performance. Overall, the model reproduced (i) the main trends across nine experimental conditions (36 predictions) for eye movement metrics, in particular the remarkable stability of the time horizon, and (ii) captured the evolution of tracking error in hand movements, although it underestimated errors at the highest scrolling velocity. This discrepancy likely arises from modeling simplifications, including the absence of signal-dependent noise, simple muscle dynamics, and limited replanning fidelity. Notably, increased information density led to higher movement frequency and greater muscle activation, whereas increased information difficulty had little effect on motor output. This suggests that energetic expenditure is more sensitive to task dynamics than to degraded sensory information, at least in the absence of risk.

Interestingly, both model and participants exhibited a “filtering effect.” With increasing information density, gaze did not strictly track high-frequency trajectory components (peaks), but instead favored smoother, lower-frequency features, resulting in increased horizontal eye–target distance without affecting minimum distance. This suggests that the visual system selectively allocates resources to task-relevant information that most improves state estimation. In contrast, increasing information gathering difficulty did not produce this filtering. Instead, gaze amplitude matched trajectory peaks, but with increased minimum distance, reflecting degraded estimation accuracy rather than selective attention.

Beyond gaze control, this paper introduced a principled formulation of hand movement based on performance optimization and energetic cost, in line with optimal feedback control models ***Todorov (2005***); ***Česonis and Franklin (2022***); ***Guigon (2023***); ***De Comite et al. (2023***). However, in contrast to standard formulations, hand planning relies on predicted future task states, yielding a feedforward component embedded within a stochastic, receding-horizon (i.e., MPC) framework. This unified formulation captures the predictive and adaptive nature of human movement, allowing replanning as new observations become available ***Takagi et al. (2025***).

Our formulation is conceptually related to active inference ***Jacques (2023***), where behavior minimizes prediction error based on hierarchical neural structures ***Rao and Ballard (1999***); ***Friston (2018***). However, rather than relying on variational free energy minimization, we adopted an MPC framework, which provides a direct and flexible formulation for motion planning and naturally accommodates reward or vigor modulation ***Ahmed and A. (2020***).

The proposed active vision framework is general and can be extended to a wide range of sensorimotor tasks. More broadly, it could be applied beyond vision to other forms of active sensing, such as exploratory touch ***Lederman and Klatzky (1993***); ***Ryan et al. (2021***), head movements for auditory localization ***Gessa et al. (2022***); ***Carlini et al. (2024***), or whole-body actions to estimate object properties and external forces ***Maurer et al. (2006***). These behaviors share a common principle: organisms actively move their sensors to optimize information acquisition for task execution. Extending the present framework to such modalities may help uncover unified computational principles governing the coordination of perception and action.

Finally, the framework offers promising technical applications. First, it could be used to predict human intention from gaze patterns, as it generates realistic eye movements grounded in optimal control principles. This could support various human–robot interaction applications, such as shared driving or robot-assisted surgery. Second, the underlying principle of active information acquisition could be transferred to artificial systems with simple adaptations of the sensorimotor parameters. Robots, drones, and autonomous agents often need to actively explore environments to gather task-relevant information ***Amigoni and Caglioti (2010***); ***Ashour et al. (2020***). Similarly, in haptics, robotic manipulators equipped with dense tactile sensors could exploit active sensing strategies to efficiently infer object properties ***Zito et al. (2013***); ***Seminara et al. (2019***); ***Dutta et al. (2026***). In both cases, the integration of perception and action through principled information-seeking control may significantly enhance performance and robustness.

## Methods

### Experiment

The trajectory tracking experiment protocol was approved by the Science, Engineering & Technology Research Ethics Committee of Imperial College London, UK (SETREC No. 21IC6578). 17 participants, aged 23-42 years old (9 males, 8 females) without known sensorimotor impairment were informed about the experiment and signed a consent form prior to starting it. Their anthropomorphic data were: height 170.3 ± 10.7 cm, weight 66.8 ± 29.2 kg and hand length 17.8 ± 5.2 cm. Each participant sat in front of a monitor (***Figure 2***A), with their right arm attached to a wrist robotic interface.

The HRX *robotic interface* (Human RobotiX, UK) was used to record wrist flexion/extension movements. The wrist joint rotation was measured with a resolution of 3.5 ⋅ 10^−6^ rad and affinely mapped to the horizontal movement of a *cursor* on the screen (motion capped at 30^°^), calibrated around a comfortable neutral hand posture. Data were recorded at 120 Hz.

*Muscle electromyography* (EMG) from the flexor carpi radialis and extensor carpi radialis longus was recorded at 2 kHz using the Wavex MiniX system (Cometa Systems). For *maximum voluntary contraction* (MVC) recording, the wrist interface was locked in a neutral position and participants were instructed to flex or extend with their maximal force for 5 s. The raw EMG signals were: (i) band-pass filtered between 50 Hz and 400 Hz, (ii) centered and rectified, (iii) low-pass Butterworth filtered with a cut-off frequency of 5 Hz, and (iv) normalized to [0,1], by dividing by the maximum post-processed activity recorded during MVC. The total activation is computed as the sum of the normalized flexor and extensor activities.

*Eye gaze* was recorded at 120 Hz using the Tobii Pro Fusion system (Tobii), following a standard five-point calibration during which participants fixed the screen center, then its four corners. The collected data were pre-processed to ensure its quality. Trials with more than 15% loss rate were discarded. Two participants (out of 17) with more than 9 invalid trials (out of 38) were excluded from the dataset. For the remaining trials, intermittent data due to blinking, large gaze deviation, or posture was handled as follows: small gaps (< 83 ms) were linearly interpolated, while larger gaps were marked as invalid. Data segments with high local velocity standard deviation (computed over a 400 ms sliding window) velocity standard deviation were classified as jittery and removed. Processed gaze data were converted from screen coordinate to eye movement angles. Saccades and smooth pursuit events were identified via peak detection on the velocity profile, using thresholds of minimum amplitude 100^°^/*s*, prominence 70^°^/*s* and distance 125 ms. The peak cut-off was set to 50% of peak height for experiment data and 100 Hz simulated data, and 30% for 1000 Hz simulated data.

We analyze eye gaze as follows. As the future target trajectory is shown to the participants on a rolling basis on the monitor, the gaze vertical position (*V-pos*) indicates the range of their planning horizon. The gaze always lead ahead the hand movement, and this range can be evaluated by dividing the gaze V-pos by the scrolling velocity (*lead time*). Combined with the gaze horizontal position (*H-pos*), the gaze trajectory can be projected onto the horizontal position-time plane (*P-pos*) for comparison with the reference trajectory and the hand trajectory. The gaze does not always land on the target trajectory. Such deviation are studied in terms of the horizontal gaze-target distance (*H-dist*) and the minimal gaze-target distance on the screen (*Dist*). An illustration of these metrics are shown in ***Figure 6***.

**Figure 6.**
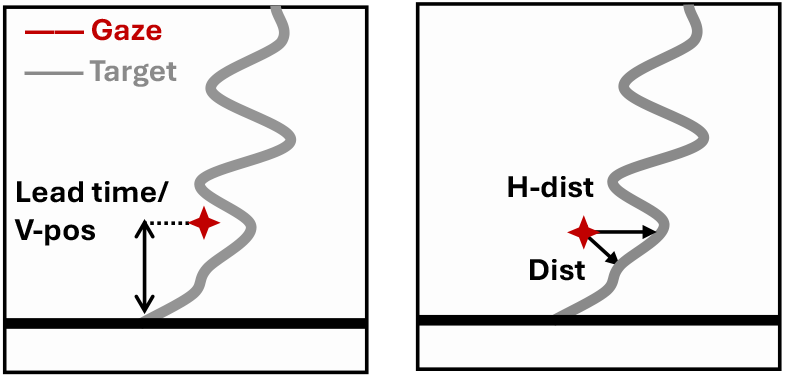
Eye movement metrics. These metrics summarize how eye gaze patterns adapt across different information-gathering conditions. They reveal, in particular, a consistent temporal lead, a “filtering” effect under specific conditions, and variations reflecting the effort cost of information acquisition.

*The task* consisted of tracking a target trajectory

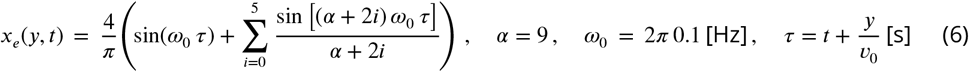

scrolling downward at constant velocity *v*_0_ = 13.4 cm/s. The displayed range, or *viewing horizon*, was *L*_0_ = 26.8 cm, corresponding to a *total time horizon T* = *L*_0_/*v* = 2 s. Participants tracked the horizontal position of the target trajectory *x*_*e*_(0, *t*) using wrist flexion/extension movements *x*_*q*_(*t*).

In the baseline condition, *L* = *L*_0_, *v* = *v*_0_, *f* = *f*_0_ (see ***Figure 2***A). The experiment began with three baseline trials. Subsequently, three experimental conditions were tested in random order (***Figure 2*** B), with the full sequence summarized in ***Figure 2*** C. Each trial consisted of 40 s of tracking task followed by 20 s of rest. Each condition included three trials. A washout trial with baseline parameters was inserted between each sequence of three trials in each condition. The order of conditions and parameter variations was pseudo-randomized across participants to mitigate learning effects. To maintain motivation, a scoreboard displaying previous performance was shown during rest periods.

*Statistical analysis* was carried out on six metrics quantifying differences in behavior across conditions {V-position, lead time, H-dist, dist, RMSE, total activation}. For each metric, normality was assessed using the Shapiro-Wilk test. Depending on the outcome, either a paired t-test (normal data) or a Wilcoxon signed-rank (non-normal data) was applied, with Bonferroni-Holm correction for multiple comparisons.

### Hand-eye control model implementation

As described in ***Equation 2*** and ***Equation 5***, eye gaze is controlled to optimize information acquisition, while the hand is tracking the target. For simplicity, we neglect proprioceptive feedback and motor noise in eye movements. Note that a complete description of the notations used throughout the paper is provided in the Appendix ***Appendix 3—table 1***. In this section, we detail the implementation of eye and wrist control strategies, where the eyes are dedicated to information gathering and the wrist to task execution. Modeling eye movements requires capturing both the increased sensitivity around the fovea and the spatiotemporal integration of sensory information.

Since *eye movements* in the experiment remain within 10^°^, we linearly map angular motion onto the screen and approximate gaze as a point mass in Cartesian coordinates with dynamics

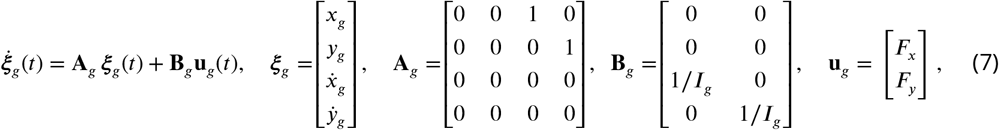

where the inertia-equivalent mass is *I*_*g*_ = 6.88⋅10^−7^ kg m^2^, derived from ocular rotational inertia (4.12⋅ 10^−7^kg m^2^) and a viewing distance of 0.6 m from eye to the screen. Viscosity is neglected as it has only a effect on effort cost, and motor noise is omitted since it primarily accounts for small corrective movements around optimal gaze positions, with negligible impact on the overall behavior studied here ***Robinson (1964***).

We model *wrist rotation*, actuated by flexor and extensor muscles, as

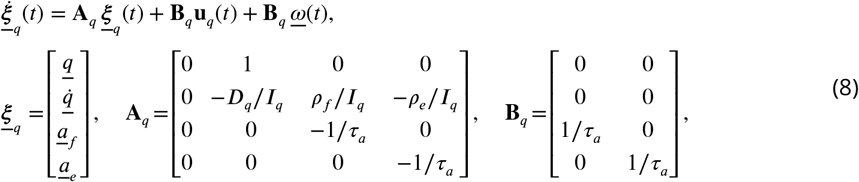

where the wrist angle *q* evolves under inertia *I*_*q*_ = 0.002 Nm⋅s^2^ and viscosity *D*_*q*_ = 0.03 Nm⋅s, taken from ***Verdel et al. (2026***). Normalized muscle excitations **u**_*q*_ = [*u*_*f*_, *u*_*e*_]^′^ ∈ [0, 1]^2^ are transformed into muscle activations [*a*_*f*_, *a*_*e*_]^′^ via first-order low-pass dynamics with time constant τ_*a*_ = 0.04 s***Gaveau et al. (2021***). Gains ρ_*f*_ = 8 Nm, ρ_*e*_ = 4.6 Nm***Yoshii et al. (2015***) map activations to torques. Motor variability is modeled as additive Gaussian noise 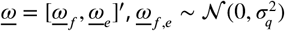 with σ_*q*_ = 0.02***Hamiltonet al.***(***2004***).

To represent the tracking task, the *target trajectory* displayed at time *t* in the (*x, y*) is modeled in screen coordinates as an infinite-dimensional vector signal

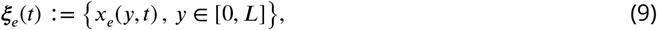

where *L* is the display height. Downward scrolling at velocity *v* induces a traveling-wave relationship

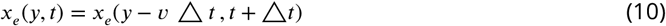

for any time interval △ _*t*_ within the range of ξ_*e*_. Visual *observations* of the target trajectory follow ***Equation 3***:

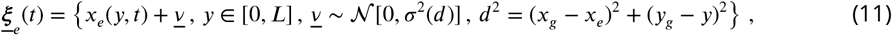

with distance dependent noise σ(*d*) described in ***Equation 3***. Similarly, the cursor (hand position) is observed as

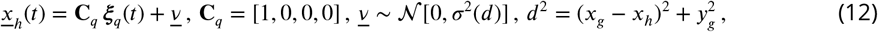

At each timestep δ*t*, estimates of the target 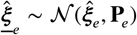 and cursor 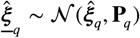are updated by integrating the new observation (<u>ξ</u>_*e*_, <u>*x*</u>_*h*_) in a stochastically optimal manner using linear quadratic estimation (LQE), with a small timestep δ*t*. The target estimate evolves via a Kalman update with spatially varying gain *K*_*e*(*y, t*), while the wrist state follows a standard discrete-time Kalman filter with process noise **Q**_*h*_. Initial conditions assume high uncertainty for the target **P**_0_ and perfect knowledge of the wrist state 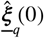. For the eye movement:

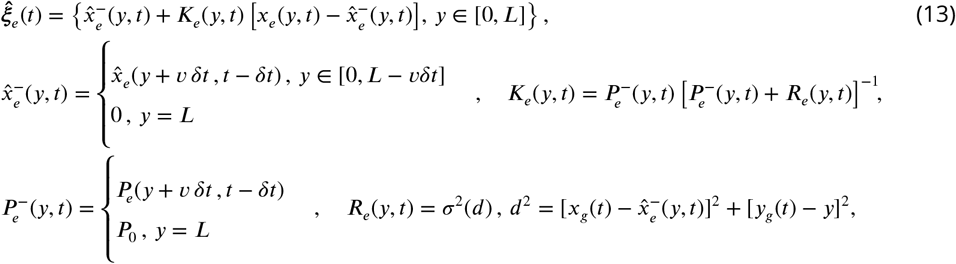

where *R*_*e*_ is the variance of observation noise at the current estimation. For the hand movement:

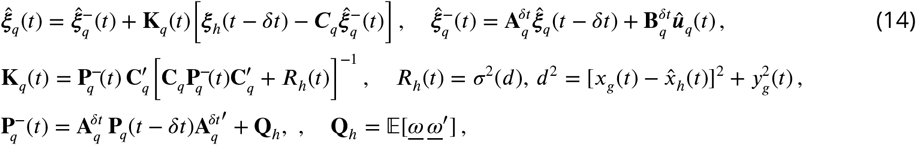

where 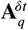, 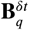 are the discrete equivalence of the joint dynamical matrices **A**_*q*_, **B**_*q*_, ω is motor noise and **Q**_*h*_ its covariance (see ***Equation 8***). The cursor position 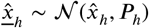 is then computed from the estimated joint state.

*Eyes’ motion* serve two objectives: (i) reducing the uncertainty of the target trajectory (*U*_*e*_ in ***Equa- tion 2***); and (ii) monitoring the cursor to estimate task errors (*U*_*h*_). Combined with control effort, this yields the eye cost function

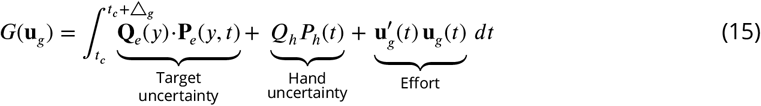

where **Q**_*e*_(*y*) = *c e*^*dy*^ = 0.0016 *e*^−68 *y*^ prioritizes information near *y* = 0, and *Q*_*h*_ = 9.2⋅10^−4^. The planning horizon △_*g*_ = 1.0 s results in an optimal control sequence 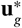, of which α_*g*_ = 67% are executed before replanning.

To control the wrist, visual information of the target trajectory is transformed from screen space to wrist joint coordinates. At time *t*_*c*_, the desired wrist trajectory over the prediction horizon is defined as

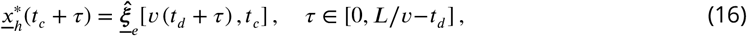

where *t*_*d*_ = 200 ms accounts for sensorimotor delay. Thus, 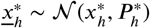 represents the predicted trajectory to be tracked from the present time *t*_*c*_ up to *L*/*v*−*t*_*d*_ into the future.

According to ***Equation 5***, Hand movements are planned to minimize tracking error and effort, yielding the cost:

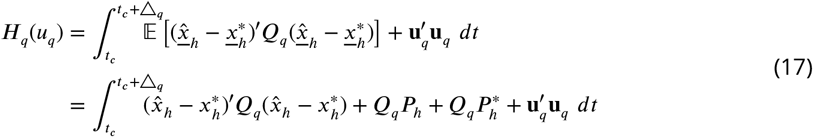

with *Q*_*q*_ = 0.11, planning horizon △_*q*_ = 0.28 s as in ***Guigon (2023***). The predicted hand state 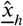 depends on **u**_*q*_, while the desired trajectory depends on the target estimate ξ_*e*_, and the two are treated as independent. At each replanning step, 45% of the optimal control sequence is executed, as in ***Guigon (2023***).

### Selection of model parameters

The model includes seven hyperparameters that are not directly available from literature. In the visual noise expression, the scaling *a* and offset *b* (***Equation 3***). In the eye cost function, the planning horizon △_*g*_ and replanning rate α_*g*_, as well as the hand uncertainty weight *Q*_*h*_ and task uncertainty **Q**_*e*_, with weight *c* and scaling *d* (***Equation 15***). In the hand cost function, there is the hand tracking error weight *Q*_*q*_ (***Equation 17***).

We performed a random search over these parameters using Latin hypercube sampling ***McKay et al. (2000***). Simulations were run at 100 Hz for 22 *s* (discarding the first 2 *s* for filter convergence) across the 9 experimental conditions. Model outputs were compared to human data across 6 metrics that characterizes their motion pattern: **gaze lead time, H-dist, saccade frequency, pursuit duration, hand RMSE, total activation**. Note that this is different from the metrics in Section Model validation as they were selected to quantitatively represent the effect of different experiment conditions, whereas here we would like to evaluate how well does a simulated trajectory resembles the average human motion pattern. The overall fitting score is defined as

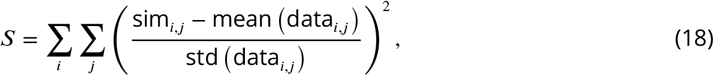

where *i* ∈ {1, 2, …, 9} refers to the experimental conditions tested in the experiment and *j* ∈ {1, 2, …, 6} are the performance metrics.

Such score evaluates the difference between the simulation value and the average participant value, scaled by the standard deviation for normalization. The best hyperparameter set is selected to produce the results in this paper and their values reported above correspondingly. A further sensitivity analysis confirmed that the model behavior is relatively stable around the selected hyperparameters (Appendix ***Appendix 2—figure 1*** -5).

## Acknowledgments

This study was funded by the European Research Council (ERC Synergy, “Natural BionicS”, grant number: 810346), and by the European Commission (FETOPEN H2020, “NIMA”, grant number: 899626 & Leadership in enabling and industrial technologies - Information and Communication Technologies (ICT), “ReHyb”, grant number: 871767).

## Gaze filtering effect with frequency

As observed both in model prediction and experiment, the eye gaze do not directly land on the trajectory; it tends to more frequently land on the inside of a curve, resulting in what is similar to a ’low-pass filtering’ effect. At lower frequencies, the gaze more closely follows the shape of the target trajectory while at higher frequencies it omits the high frequency components, as shown in n ***Appendix 1—figure 1***.

**Appendix 1—figure 1.**
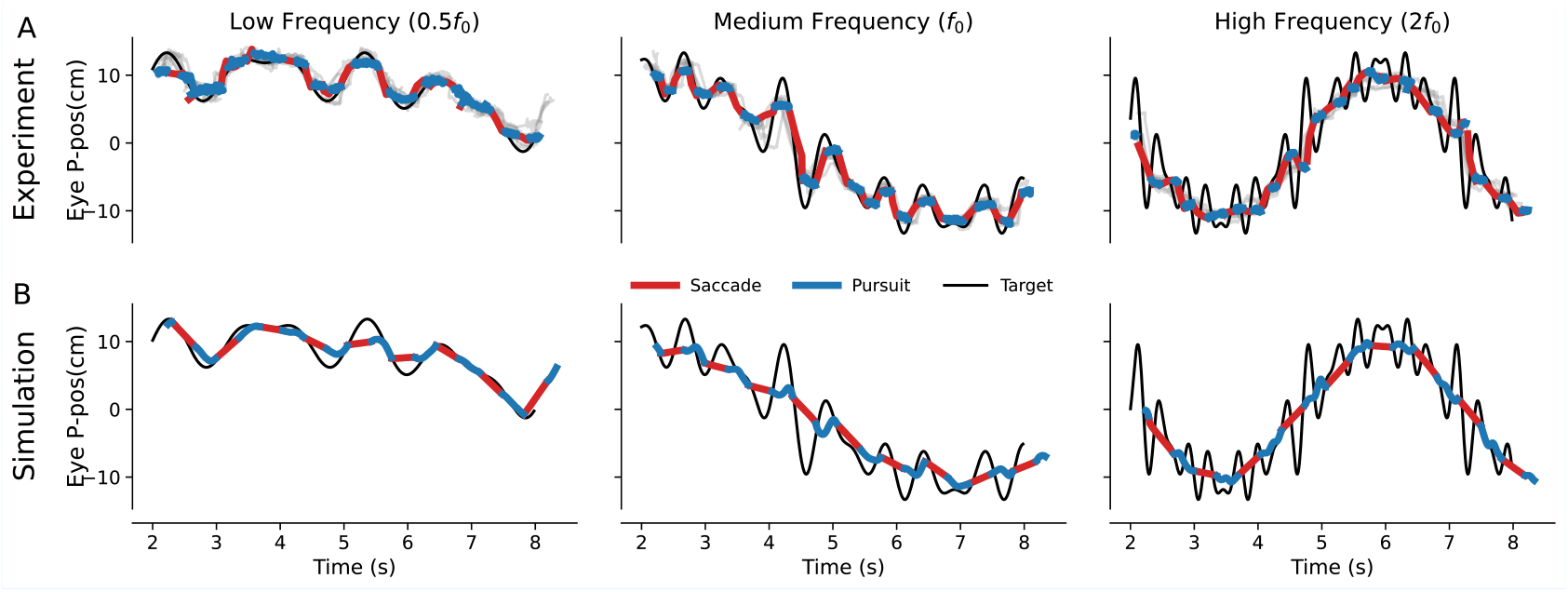
Gaze filtering filtering of high-frequency eye movement. Comparison of the movement trajectories projected onto the ne (P-pos) across different target trajectory frequencies. In all panels, gaze is segmented into saccades (red) mpared to the target trajectory (black). **A:** Experiment data for a representative participant, highlighting one in gray. **B:** Model output for a selected parameter set, sampled at 100 Hz

### Sensitivity analysis

In the main text of the manuscript, we presented the simulation result with the one selected hyperparameter set. Here we preform a sensitivity analysis, in which each hyperparameter is perturbed around its optimal value. The eye planning horizon and replan rate is assumed to model the model performance on a linear scale while the others on an exponential scale. The linear parameters are perturbed within ±20% of their optimal value, with a 2% step size. The exponential parameters are perturbed by [1/1.5, 1/1.48, …, 1/1.02, 1, 1.02, …, 1.48, 1.5] × original value.

A simulation of the perturbed parameter is run for each of the 8 experimental condition for 22s, with the first 2s discarded. The simulation result is evaluated using the same metrics as before: V-position, Lead time, H-dist, Dist, RMSE and total activation.

**Appendix 2—figure 1.**
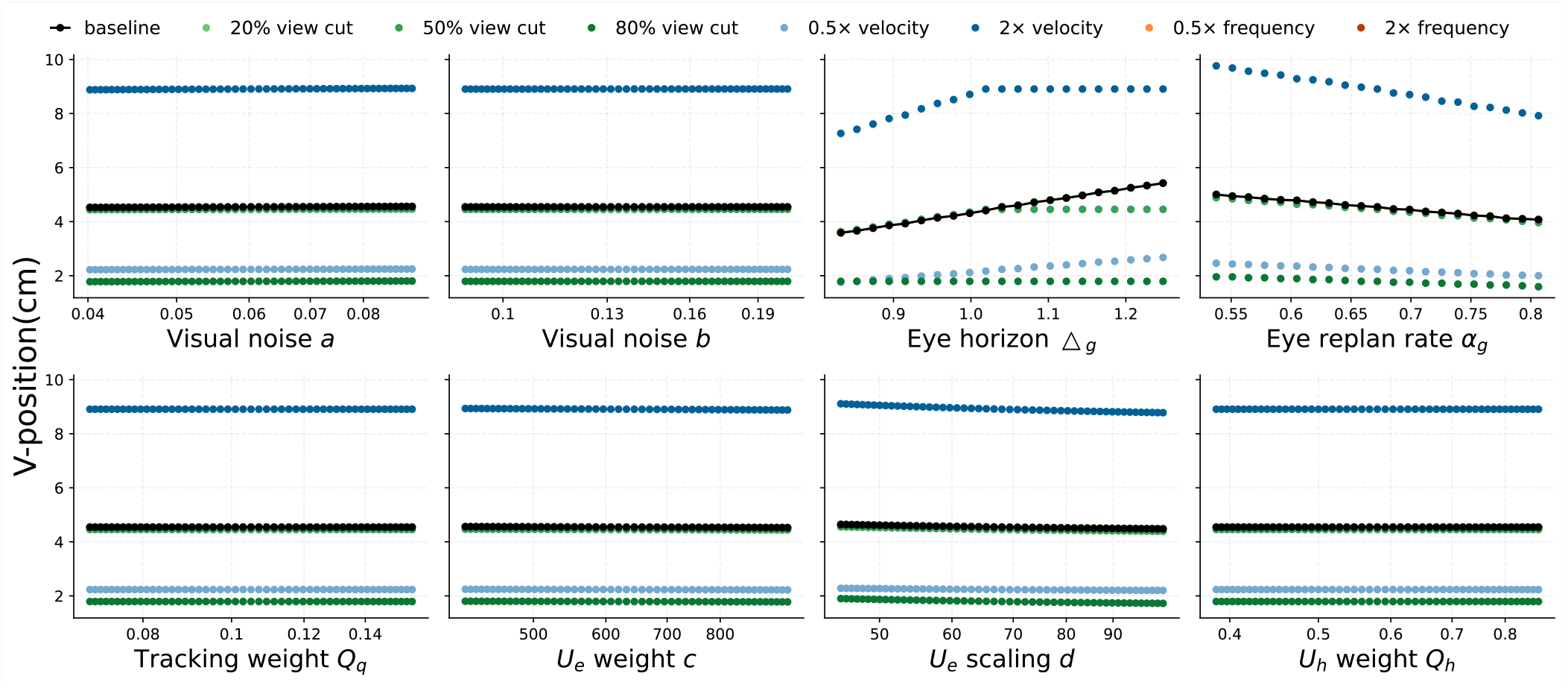
Sensitivity analysis: V position of the eye gaze

**Appendix 2—figure 2.**
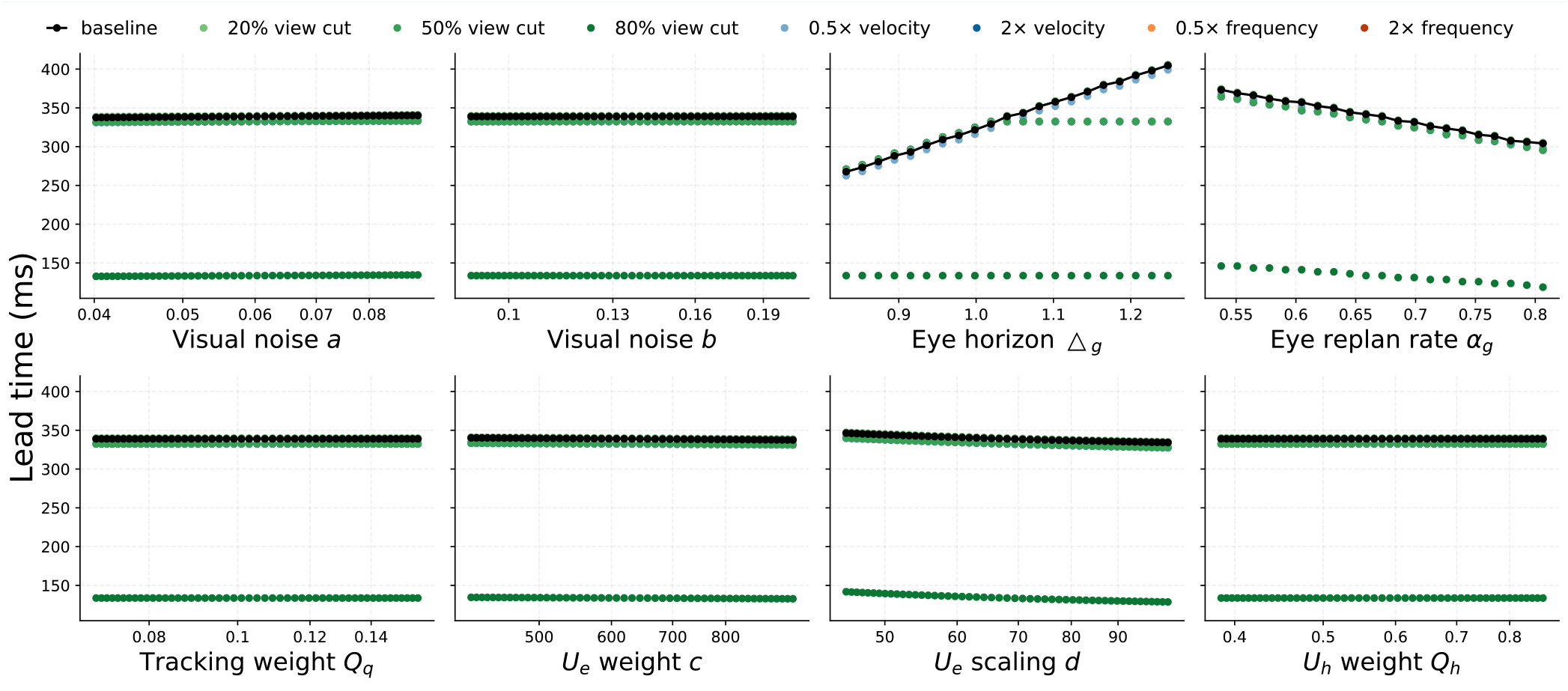
Sensitivity analysis: Lead time of the eye gaze The sensitivity analysis result shows that, the V-position and lead time increases with △_*g*_ and decreases with α_*g*_ ; it also slightly decreases with *d* (***Appendix 2—figure 1, Appendix 2—figure 2***)

**Appendix 2—figure 3.**
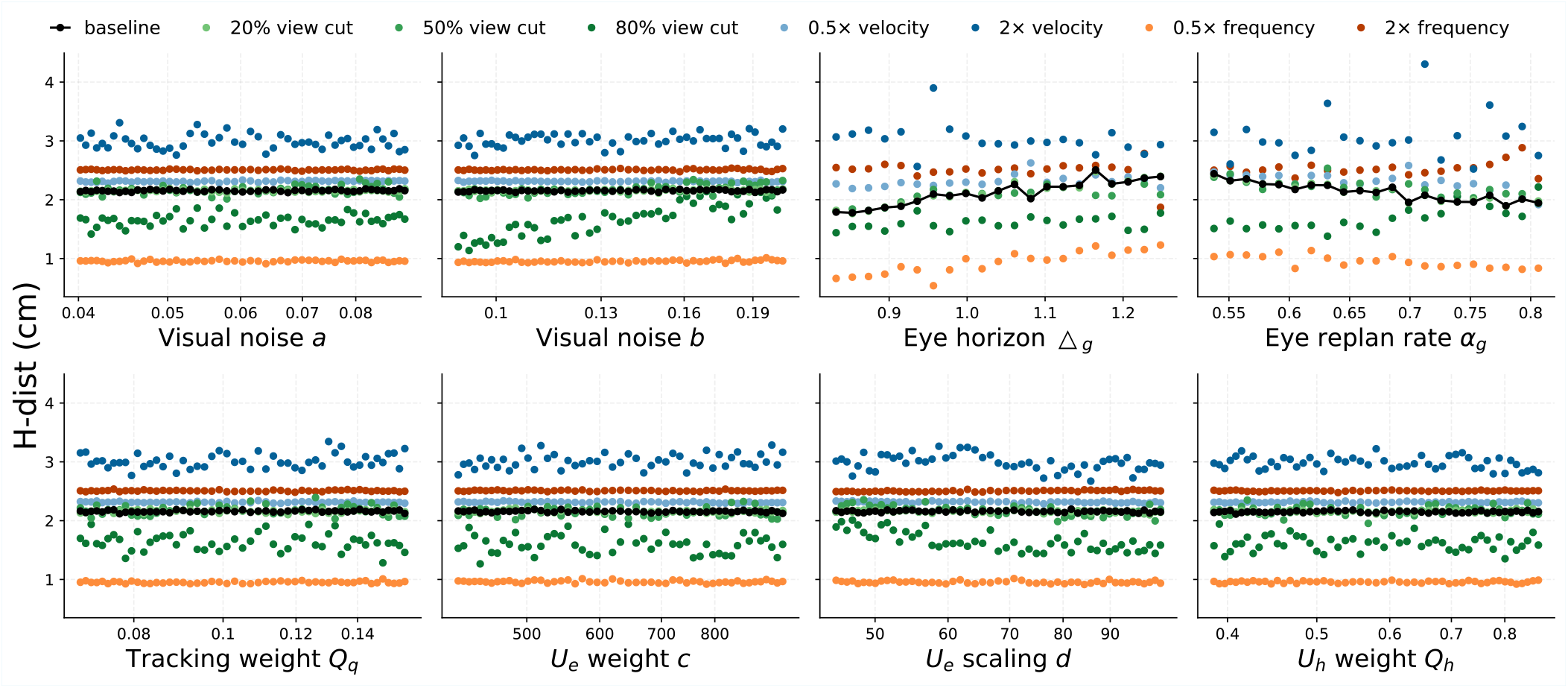
Sensitivity analysis: horizontal distance between the eye gaze and the target trajectory

**Appendix 2—figure 4.**
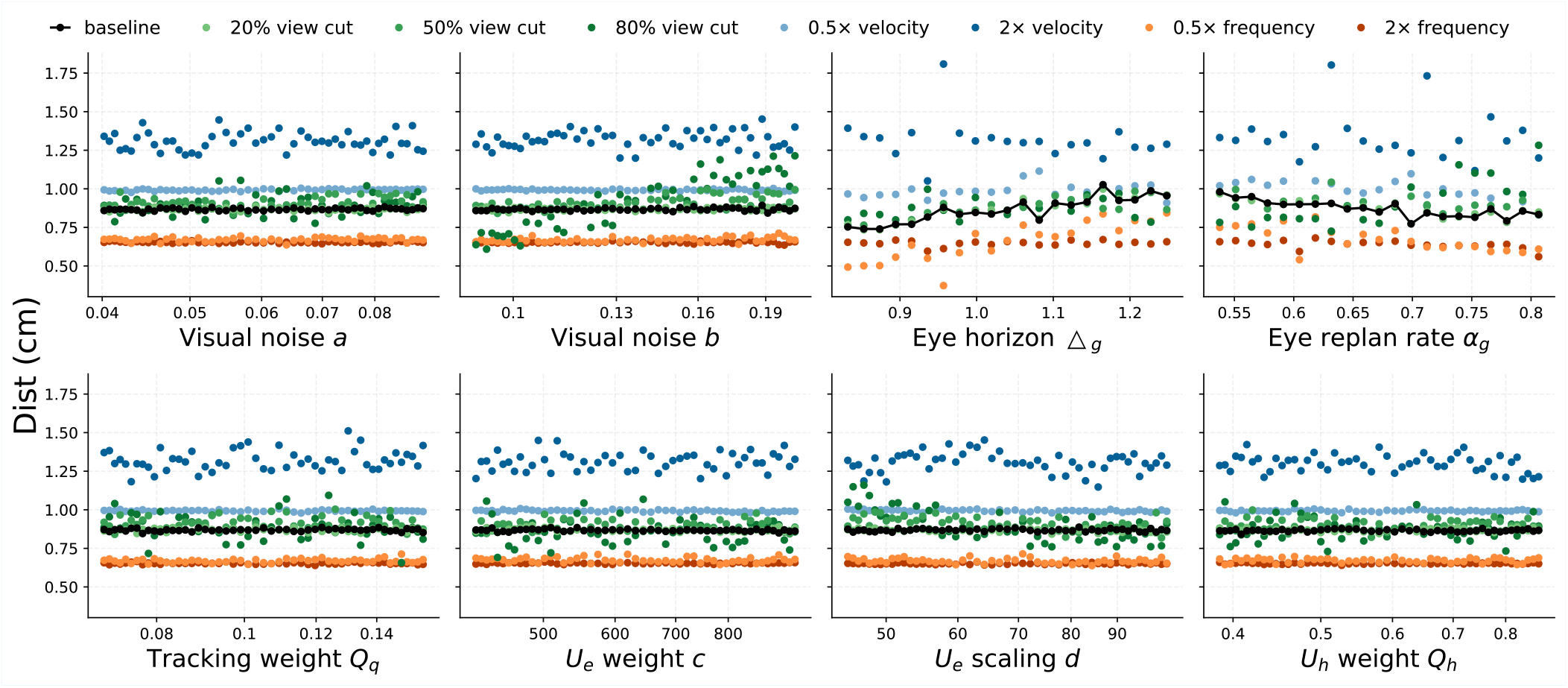
Sensitivity analysis: 2-dimensional distance between the eye gaze and the target trajectory The horizontal and absolute distance between the gaze and target increases with △*g* and decreases with α_*g*_ . Despite no obvious change for the baseline condition, the H-distance and Distance both increased with the visual noise offset at the 80% view cut condition, potentially due to reduced target trajectory estimation accuracy. (***Appendix 2—figure 3, Appendix 2—figure 4***) .

**Appendix 2—figure 5.**
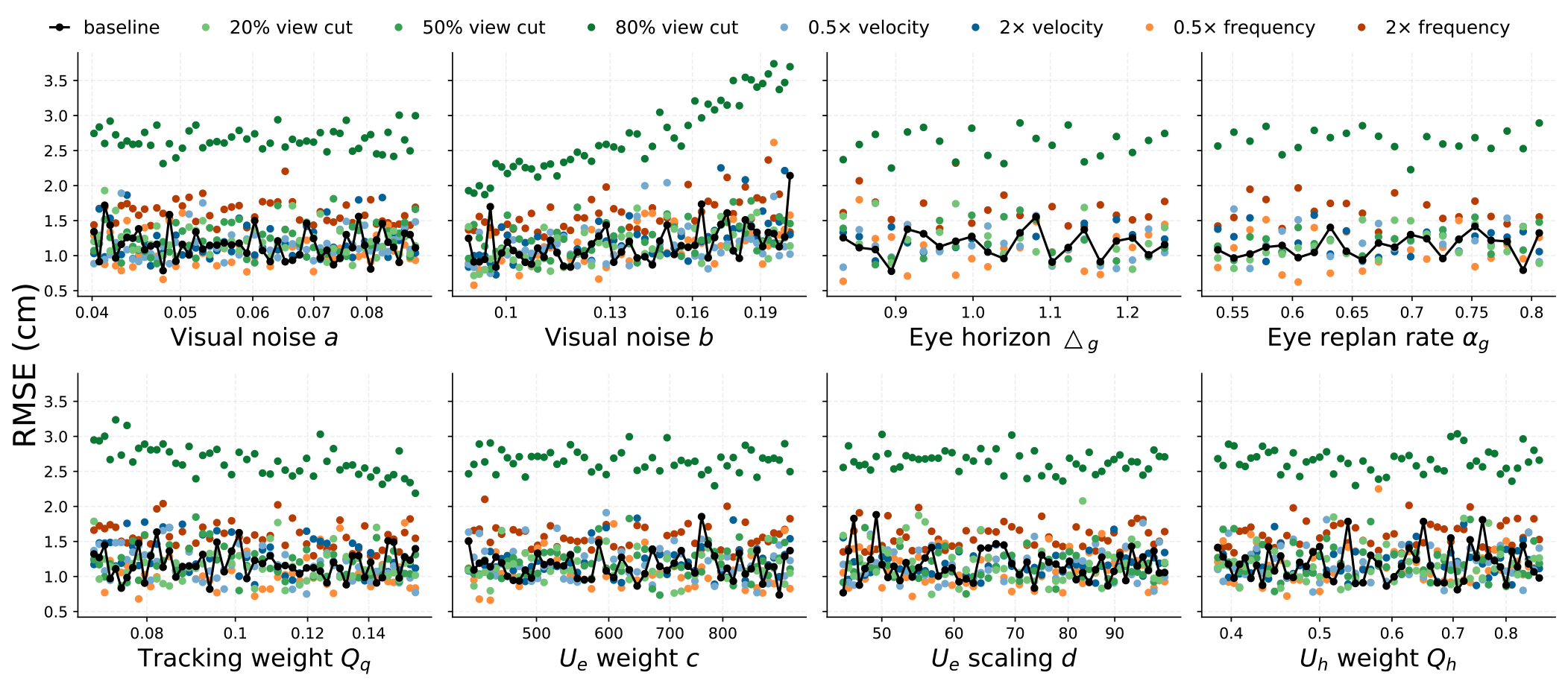
Sensitivity analysis: root mean square of the hand tracking error No significant trend can be observed for the RMSE in the baseline condition, however, with 80% view cut, RMSE is increased by higher visual noise offset, potentially due to decreased target estimation quality and. RMSE is decreased by higher hand tracking error weight *Q*_*q*_, potentially due to higher penalty on tracking error over effort. (***Appendix 2—figure 5***)

**Appendix 3—table 1.**
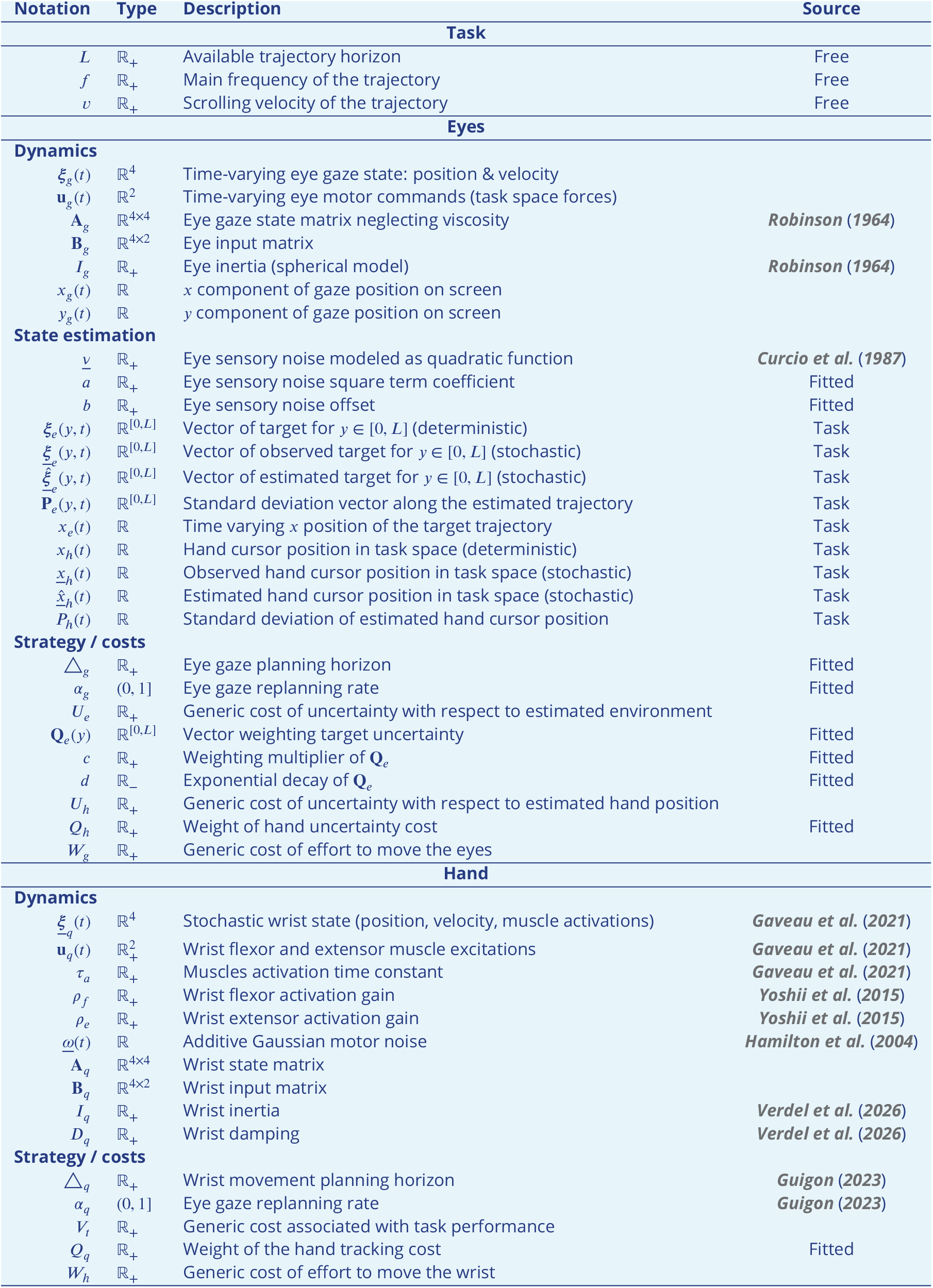
Summary of notations.

